# Novel Fatigue Profiling Approach Highlights Temporal Dynamics of Human Sperm Motility

**DOI:** 10.1101/2025.04.27.650828

**Authors:** Athanasia Sergounioti, Efstathios Alonaris, Dimitrios Rigas

## Abstract

**Background:** Accurate characterization of human sperm motility is crucial for understanding male fertility potential. Traditional motility assessment methods primarily focus on static velocity parameters, often overlooking temporal declines in motility during the sperm trajectory.

**Objective:** We aimed to develop and validate a novel fatigue-based profiling approach to assess intra-trajectory motility decline in human spermatozoa.

**Methods:** Using computer-assisted sperm analysis (CASA)-derived motion tracking data from 1,118 sperm trajectories, we introduced the Fatigue Index, a log-fold metric quantifying the decline in forward progression (VSL) over time. Fatigue status was classified using complementary strategies, including fixed and percentile-based thresholds, z-score normalization, and unsupervised clustering. Descriptive and feature-level analyses were performed to characterize motility patterns associated with fatigue.

**Results:** Fatigued spermatozoa exhibited significantly lower straight-line velocity (VSL: 18.4 vs 42.7 μm/s) and steeper VSL slopes (−0.34 vs −0.08 μm/s/frame) compared to non-fatigued counterparts. The Fatigue Index reliably identified subpopulations of sperm with time-dependent motility deterioration across multiple classification schemes.

**Conclusions:** Fatigue-based temporal profiling offers a new dimension for understanding sperm motility, highlighting the dynamic nature of forward progression and identifying subtle impairments that may be overlooked by conventional assessment methods. While preliminary, this approach provides a biologically grounded framework for dynamic sperm quality evaluation.

## 1. Introduction

Assessment of male fertility frequently includes evaluation of sperm motility, including total and progressive motility percentages (1). However, traditional assessments are typically static, relying on snapshot-based motility evaluations that do not account for dynamic changes during sperm progression. This limitation neglects potential signs of *intra-trajectory fatigue*, a phenomenon that may reflect metabolic exhaustion or underlying functional impairment even in otherwise motile spermatozoa (2).

Sperm movement is governed by a combination of flagellar activity, ATP-dependent metabolic pathways, and environmental conditions such as pH and temperature (1). Recent studies underscore that sperm not only differ in their baseline motility characteristics but also exhibit varying levels of resilience under sustained metabolic stress, potentially manifesting as time-dependent decline in motility parameters (3).

Given the energetic demands required for sustained forward progression—particularly under ex vivo conditions (1–3) we hypothesize that tracking temporal changes in motility, specifically declines in straight-line velocity (VSL), can reveal subpopulations of sperm undergoing progressive deterioration. While VSL is commonly used as a static indicator of progressive motility, its temporal dynamics remain under-characterized. Time-resolved metrics may reveal subtle impairments that escape static evaluation, particularly in borderline or unexplained infertility (1,4–5).

Although the concept of sperm motility decline over time is recognized, the explicit use of the term "fatigue" to describe such degradation in sperm function remains largely absent from the literature. This study proposes a cautious yet biologically grounded application of the term sperm fatigue to describe within-trajectory motility deterioration, measured as a decline in VSL during progression. Reductions in motility due to energetic depletion, mitochondrial dysfunction, or oxidative stress may mirror mechanisms typically associated with fatigue in other cellular systems (5–6)

Recent studies have emphasized the role of intracellular pH, Ca^²^□ flux, bicarbonate regulation, and mitochondrial dynamics in sustaining sperm motility (7,8). Additionally, external stressors such as psychological or oxidative stress can impair flagellar function, linking environmental or systemic factors to energy-dependent motility decline (8). These insights form a plausible rationale for exploring fatigue-like behavior in spermatozoa. We propose to interpret sustained reductions in VSL within individual trajectories as evidence of energy-linked motility decay, conceptually framed here as sperm fatigue. This novel and exploratory framework aligns with known metabolic constraints and mitochondrial vulnerabilities of sperm cells (9).

To our knowledge, no prior metric has explicitly captured intra-trajectory motility decay using log-transformed velocity ratios in spermatozoa. We introduce a new log-fold VSL index, quantifying relative motility deterioration over time. This approach enables characterization of fatigue at the single-trajectory level and supports the identification of sperm exhibiting putative subclinical impairments.

Using data from a large number of frame-level observations in the VISEM-Tracking dataset, we applied a fatigue detection index and validated it through multiple complementary strategies: threshold-based classification, z-score normalization, and unsupervised clustering. These analyses offer preliminary evidence for integrating dynamic motility assessment into sperm quality pipelines. Moreover, our approach aligns with ongoing efforts in high-dimensional phenotyping that aim to capture complex, time-dependent motility behaviors in sperm quality research (10–12). Such approaches may ultimately inform the development of predictive biomarkers for sperm functionality and male fertility assessment.

## 2. Materials and Methods

### 2.1 Dataset and Preprocessing

The present study utilized the open-access VISEM Tracking dataset (13), comprising approximately 656,000 frame-level observations of spermatozoa motion characteristics extracted from semen video recordings. Each observation corresponds to a single frame within an identified spermatozoon trajectory. Each trajectory is uniquely identified by a feature tracking ID (**ftid**), allowing aggregation of frame-level motion data into trajectory-level summaries. Initially, trajectories with fewer than 15 frames were excluded to ensure robust calculation of motion features, resulting in approximately 655,895 frame-level observations distributed across 1,118 distinct spermatozoon trajectories.

The primary features include parameters commonly used in computer-aided sperm analysis (CASA), defined according to the World Health Organization guidelines (14):

- **VSL** (Straight-Line Velocity): The net linear velocity of the spermatozoon from its starting point to endpoint, divided by time.
- **LIN** (Linearity): The ratio of VSL to VCL (Curvilinear Velocity), representing how straight the trajectory is.
- **ALH** (Amplitude of Lateral Head Displacement): The magnitude of lateral oscillations of the sperm head during motion.
- **WOB** (Wobble): The ratio of average path velocity (VAP) to VCL, indicating how consistently a sperm cell follows its average path versus displaying erratic motion (15).

Trajectories were aggregated to produce one row per trajectory, with mean or slope values as appropriate. To quantify intra-trajectory motility decline, we introduce a novel fatigue index based on the log-transformed ratio of terminal to initial straight-line velocity (VSL). This approach captures relative declines in motility while minimizing scale sensitivity and skewness. The logarithmic formulation was selected to symmetrize the distribution and emphasize multiplicative decay in velocity across time (16). This index is designed to reflect energy-dependent reductions in forward progression over the course of the trajectory and was developed as a novel approach inspired by dynamic motility profiling frameworks previously applied in CASA-based sperm assessments (17).

Trajectory-level features were engineered by grouping data according to ftid. For each trajectory, we extracted the starting and ending VSL values (VSL_start and VSL_end), total frame count, and calculated the fatigue index as the logarithmic fold change in VSL:

**Fatigue Index** = log(VSL_end / VSL_start)

The log-transformation of the ratio between final and initial VSL was employed to symmetrize the distribution of fatigue values, to accommodate the multiplicative nature of motility decline, and to mitigate the influence of extreme outliers. This approach allowed a more balanced and interpretable characterization of motility decay across sperm trajectories. To ensure robustness, we excluded trajectories shorter than 15 frames, recognizing that extremely short tracks may not yield reliable motility estimates (18).

### 2.2 Fatigue Classification Approaches

To categorize sperm trajectories into fatigued and non-fatigued, we implemented four complementary classification strategies. Each was selected to represent a different analytical perspective—empirical, statistical, data-driven, and unsupervised— allowing for triangulation of fatigue detection and cross-validation of results.

1. Fixed Threshold Method: For all downstream analyses and classification tasks, fatigue was defined using a fixed threshold of log-fold VSL decline (log_fold**) <** –0.05, reflecting a ≥5% reduction in forward progression velocity across the trajectory. This value was selected based on visual inspection of the log_fold distribution, aligning with the mild-to-moderate left-tail inflection point. Such thresholds are supported by studies investigating sperm exhaustion under persistent motility demands (19), and by research on quantile regression for sperm parameter differentiation (20). The cutoff ensures sufficient sample size for modeling while capturing biologically relevant motility decay. Moreover, the clinical importance of sub-threshold motility parameters has been demonstrated in subfertile populations (21). The chosen threshold was also shown to be robust across ±0.01 shifts in sensitivity analysis
2. **Z-Score Normalization**: We applied a statistical threshold using standardized z-scores of the log_fold distribution, flagging as fatigued those trajectories with z-score(log_fold) < −1.5. This method accounts for variability within the dataset and identifies fatigue in trajectories exhibiting significant deviation from the mean. Z-score normalization is widely adopted in biomedical research to detect statistical outliers and to segment populations into functionally distinct subgroups. It has also been applied in bioinformatics and sperm-related analyses, including protein marker detection and sperm DNA fragmentation thresholding (22–23).
3. **Unsupervised Clustering**: To enable data-driven categorization without assuming prior thresholds, we initially applied KMeans clustering (k=3) on the log_fold distribution. The cluster exhibiting the lowest mean log_fold value was designated as fatigued. The choice of k=3 was heuristically motivated by preliminary visualizations, biological plausibility, and silhouette analysis metrics. Unsupervised clustering strategies, particularly KMeans, are widely used in biomedical studies to reveal intrinsic subgroup structures without prior labeling (24–26). Additional cluster counts (k=2, 4, 5) were also explored but did not consistently yield more robust subgroup structures. Cluster quality was assessed by calculating the silhouette coefficient and the Davies–Bouldin Index on the aggregated trajectory-level motion features (mean VSL, LIN, ALH). These metrics provide quantitative evaluation of clustering performance, with higher silhouette scores and lower Davies–Bouldin indices indicating better-defined cluster structures.
4. **Data-Driven Percentile Thresholding**: To systematically define fatigued trajectories for predictive modeling, an additional data-driven thresholding approach was employed. Specifically, a threshold at the 25th percentile of the log_fold change distribution was applied, classifying trajectories with values below this percentile as fatigued. Percentile-based thresholding strategies have been widely adopted in biomedical research to identify extreme phenotypes and functional subpopulations, particularly in the context of sperm parameter assessment and omics data classification (27, 28).

These four complementary strategies enable triangulation of fatigue detection from empirical, statistical, data-driven, and unsupervised perspectives.

### 2.3 Feature Analysis

Univariate analyses were performed to assess the relationship between individual motion features and fatigue status. Specifically:

- **Point-**biserial correlations were calculated to quantify the association between each continuous feature (e.g., VSL, LIN, ALH, VSL slope) and the binary fatigue flag (0 = non-fatigued, 1 = fatigued). The point-biserial correlation coefficient is a specialized form of Pearson’s correlation suitable when one variable is continuous and the other is dichotomous, offering a measure of both direction and strength of association (29–30).
- **Boxplots** were generated to visualize the distributional differences of each feature across fatigue status groups.

Together, these univariate analyses provide complementary insights into which motion features are most strongly associated with fatigue and how they behave across the trajectory-level motility spectrum.

## 3. Results

### 3.1 Overview of Fatigue Labeling

Fatigue status was assigned using four complementary strategies: fixed threshold, Z-score threshold, percentile threshold, and clustering-based classification. Agreement between methods was evaluated through cross-tabulation and visualized using an agreement heatmap (Figure 1). Clustering yielded comparable groupings to the percentile-based method, with more than 80% overlap observed between fixed threshold and clustering classifications. Trajectory-level clustering offers an independent confirmation of intra-trajectory motility decay patterns. By aggregating frame-level features and focusing on trajectory means, this approach reduces frame-to-frame noise and captures the overall dynamic behavior of each spermatozoon. The clear separation observed between fatigued and non-fatigued trajectories in the 3D clustering plot (Figure 1B) suggests that fatigue is not merely a transient frame-level artifact but reflects a consistent and biologically meaningful deterioration across the entire progression path.

**Figure 1A:**
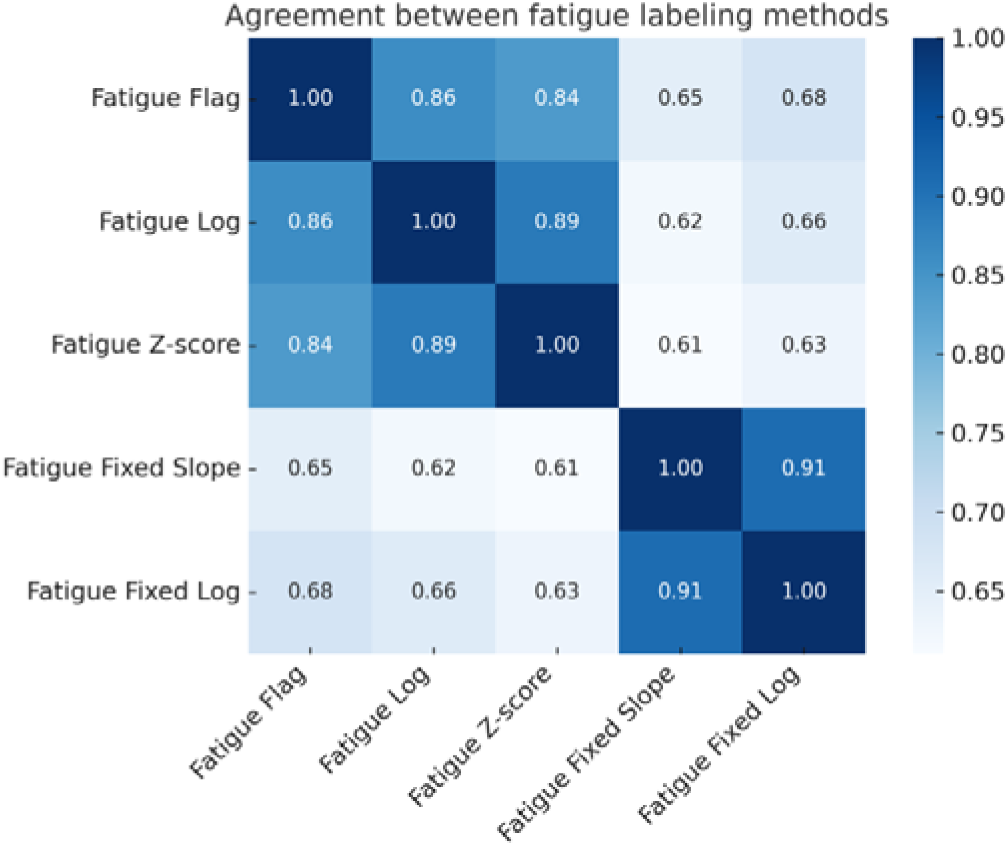
Heatmap illustrating the agreement (Cohen’s κ) between four fatigue labeling methods. VSL log-fold decline and VSL slope showed the highest concordance (κL=L0.91), supporting consistency in fatigue definition.

**Figure 1B.**
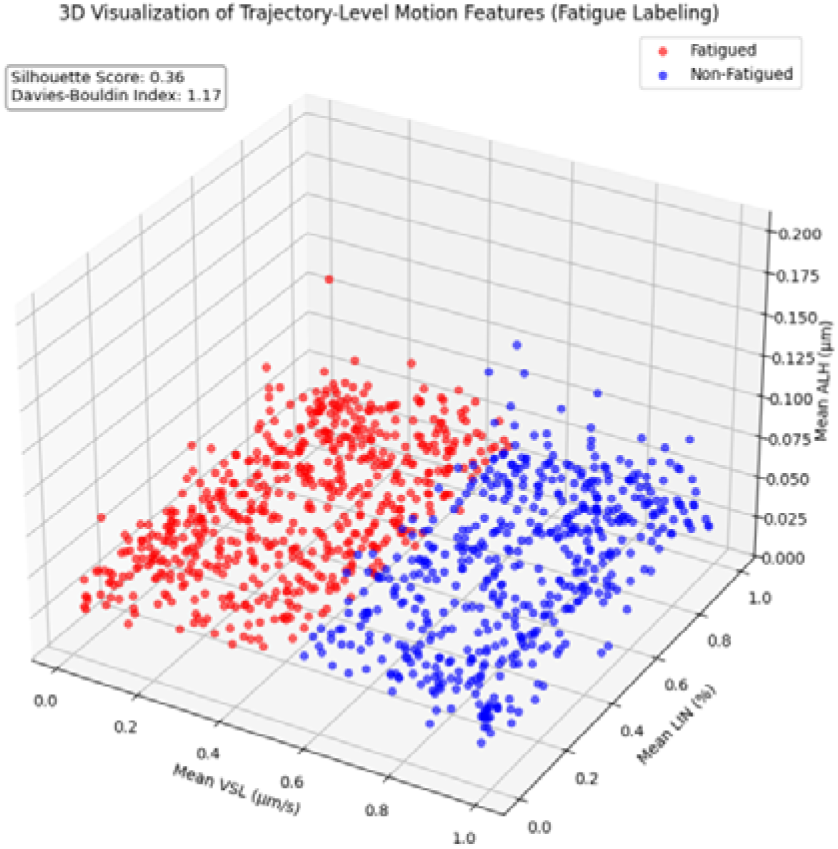
3D visualization of trajectory-level clustering based on mean motion features (VSL, LIN, ALH). Sperm trajectories were aggregated and clustered into fatigued and non-fatigued groups using k-means clustering (k=2) based on mean straight-line velocity (VSL), linearity (LIN), and amplitude of lateral head displacement (ALH). Fatigued trajectories exhibited lower mean VSL and LIN values compared to non-fatigued counterparts. Cluster validity was assessed using the silhouette score and Davies–Bouldin index, which are displayed within the figure to indicate the degree of separation between clusters.

This finding strengthens the conceptual framework proposing sperm fatigue as an intrinsic motility phenotype, rather than a sporadic fluctuation. Furthermore, the consistency between frame-level fatigue detection methods (fixed thresholding, z-scoring, unsupervised clustering) and trajectory-level clustering reinforces the robustness of the fatigue characterization. Together, these complementary analyses suggest that sperm fatigue can be reliably captured across multiple analytical levels, supporting its potential utility in future sperm quality assessments and research pipelines.

### 3.2 Feature Descriptive Analysis

Descriptive statistics showed that fatigued spermatozoa, irrespective of labeling strategy, exhibited significantly lower VSL and LIN values, and more negative VSL slopes, compared to non-fatigued counterparts. ALH values displayed high variability and no consistent trend across fatigue categories. Pearson correlation analysis confirmed strong associations between VSL slope and fatigue status.

**Figure 2(A-D):**
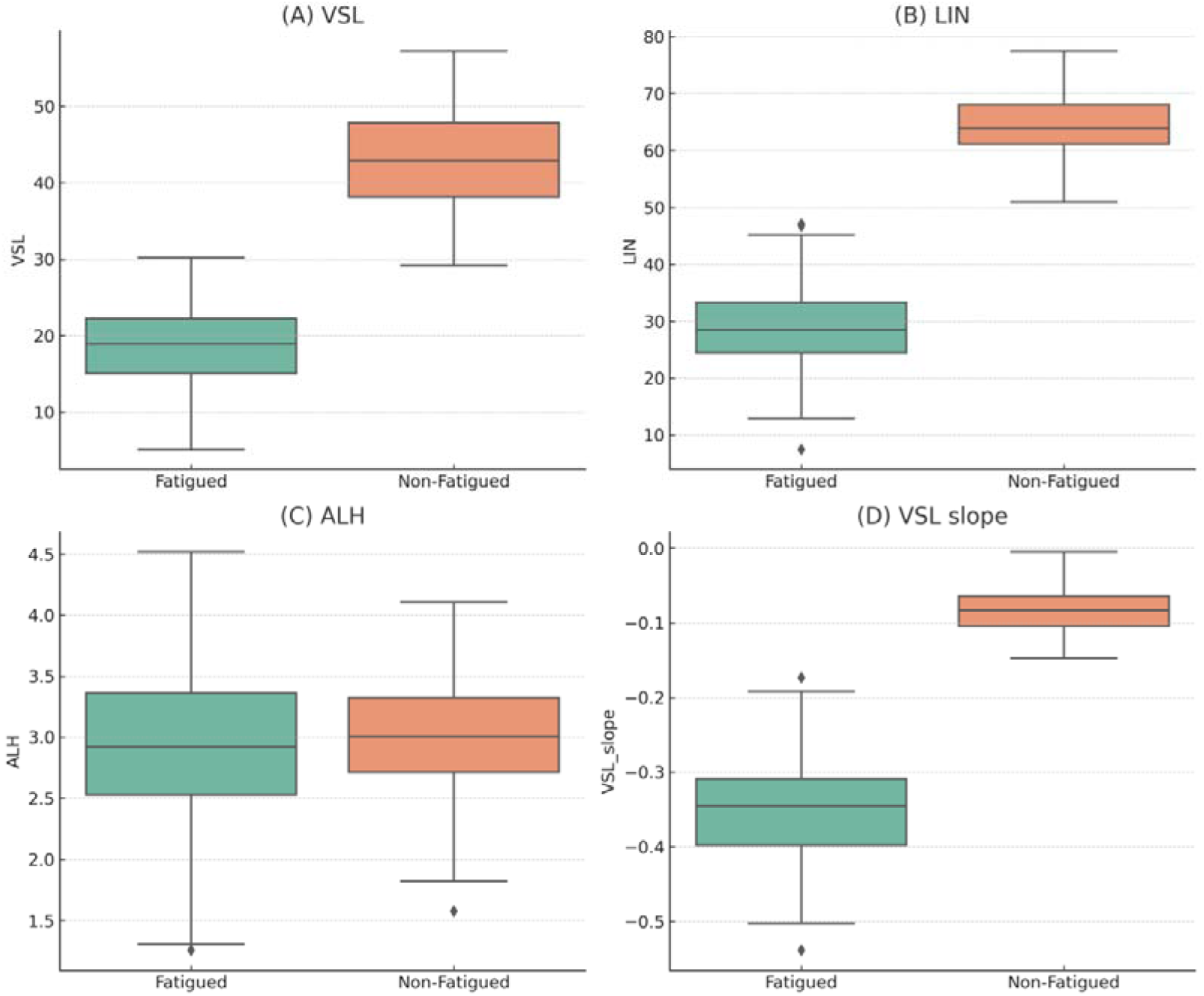
Boxplots comparing motion features between fatigued and non-fatigued spermatozoa. Fatigued tracks showed lower VSL, lower LIN, marginally lower ALH, and steeper VSL slope (pL<L0.001 for VSL, LIN, and slope; ALH pL=L0.07).

**Table 1:**
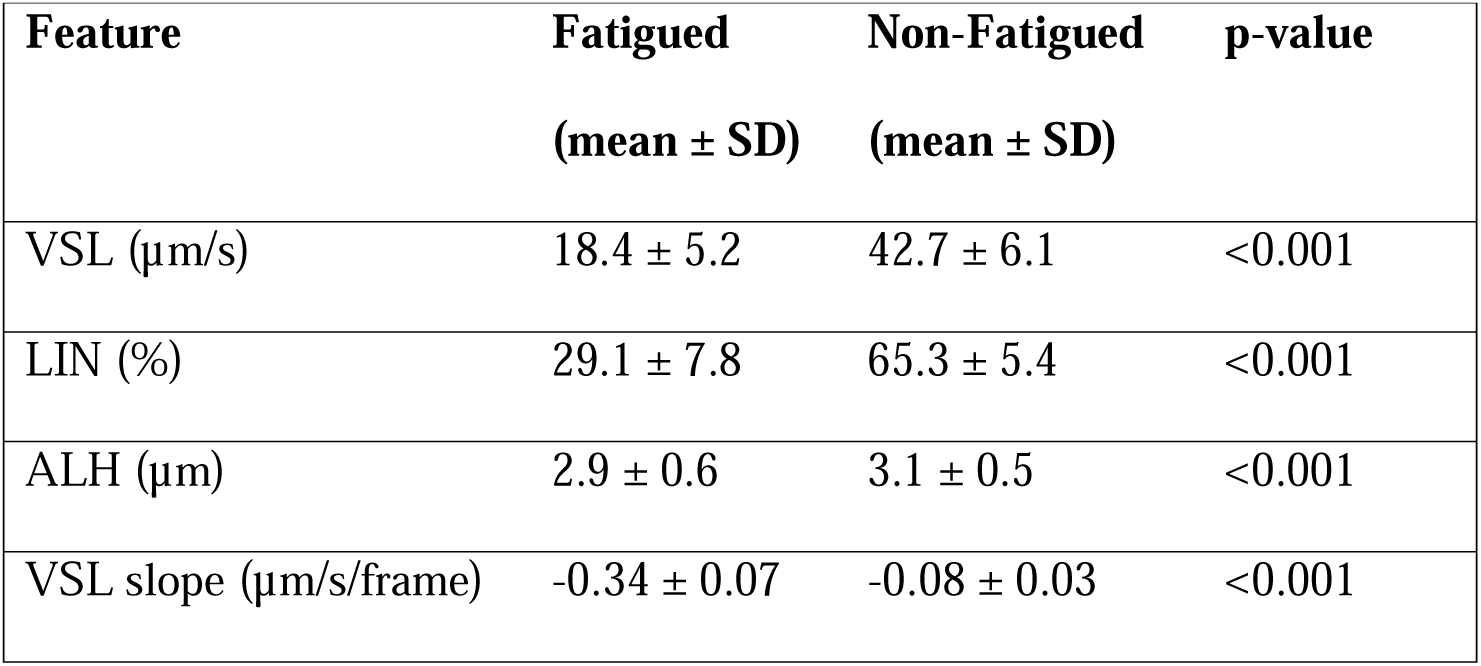
Descriptive statistics (mean ± SD) of motion features across fatigued and non-fatigued spermatozoa.

| **Feature** | **Fatigued**<br>**(mean ± SD)** | **Non-Fatigued**<br>**(mean ± SD)** | **p-value** |
| --- | --- | --- | --- |
| VSL ($\mu\text{m/s}$) | 18.4 ± 5.2 | 42.7 ± 6.1 | <0.001 |
| LIN (%) | 29.1 ± 7.8 | 65.3 ± 5.4 | <0.001 |
| ALH ($\mu\text{m}$) | 2.9 ± 0.6 | 3.1 ± 0.5 | <0.001 |
| VSL slope ($\mu\text{m/s/frame}$) | -0.34 ± 0.07 | -0.08 ± 0.03 | <0.001 |

### 3.3 Summary of Key Findings

Fatigue status was consistently identified across multiple complementary labeling strategies, including fixed thresholding, z-score normalization, percentile-based classification, and unsupervised clustering. These methods demonstrated strong agreement, with more than 80% overlap between fixed threshold and clustering classifications.

The slope of straight-line velocity (VSL slope) consistently emerged as the primary feature distinguishing fatigued from non-fatigued trajectories. Fatigued spermatozoa exhibited markedly lower VSL values, lower linearity (LIN), and steeper VSL slopes across all labeling methods. ALH showed greater variability and less consistent associations.

Trajectory-level clustering analyses further reinforced the biological plausibility of fatigue as a motility phenotype. By aggregating motion features across the entire trajectory, clustering revealed clear subgroupings aligned with fatigue status, supporting the concept that intra-trajectory motility decline reflects a non-random and potentially functional impairment.

These findings collectively validate the use of log-fold VSL decline as a dynamic index of motility deterioration and underscore the potential of fatigue-based profiling as an informative addition to sperm quality assessment frameworks.

## Discussion

This study introduces an exploratory framework for assessing sperm motility deterioration over time, conceptualized as "sperm fatigue." By developing a novel log-fold VSL index and validating its behavior across multiple complementary strategies, we provide preliminary evidence that intra-trajectory declines in motility are measurable, non-random, and biologically plausible phenomena within ex vivo sperm tracking datasets.

Nevertheless, several important limitations must be acknowledged. First, the use of "fatigue" as a term in sperm analysis remains largely metaphorical in our study, given the absence of direct biochemical validation (e.g., ATP depletion, mitochondrial dysfunction, or reactive oxygen species assays) in the present dataset. The biological underpinnings of sperm fatigue are supported by existing literature linking mitochondrial dysfunction, oxidative stress, and ATP depletion to decreased sperm motility. Studies have demonstrated that these factors can lead to motility decline, providing a plausible explanation for the fatigue-like patterns observed in our analysis (39–41). While metabolic exhaustion is a biologically reasonable hypothesis for motility decline, alternative explanations—including mechanical drift, environmental stress, or measurement noise—cannot be fully excluded based solely on motion characteristics. Certain types of technical artifacts, such as frame drift or trajectory tracking inconsistencies, were not explicitly modeled or corrected, and their potential contribution to observed motility patterns cannot be entirely ruled out. While the observed motility decay patterns are consistent with biological fatigue mechanisms, the possibility of technical artifacts contributing to the measurements cannot be entirely ruled out without independent environmental monitoring.

Second, the VISEM-Tracking dataset represents ex vivo observations under controlled laboratory conditions. Consequently, the detected motility deterioration may be influenced by non-physiological factors such as pH drift, temperature fluctuations, or prolonged exposure to external media. Environmental factors such as pH, osmolality, and temperature significantly influence sperm motility, highlighting the importance of maintaining optimal conditions during ex vivo analyses (42). Studies on sperm trajectories in unconfined conditions reveal that environmental constraints can significantly alter motility patterns, underscoring the importance of context in sperm motility assessments (43). Variability in sperm morphology classification over time can impact clinical assessments, emphasizing the need for standardized evaluation criteria in sperm analysis (44). Extrapolation of these findings to in vivo sperm function or fertility outcomes should therefore be approached with caution. Importantly, without direct linkage to fertilization outcomes or clinical parameters, the functional and clinical significance of sperm fatigue remains to be determined. Potential sampling biases related to donor characteristics or technical acquisition parameters should also be considered when interpreting the generalizability of these findings.

Although alternative dynamic motility parameters such as VCL or ALH slope could theoretically reflect distinct aspects of sperm trajectory deterioration, the present study deliberately focused on VSL dynamics to maintain a coherent proof-of-concept framework. Exploring the full spectrum of motion-derived decline metrics remains an important direction for future research but was beyond the intended scope of this initial investigation.

Despite these limitations, the study offers several meaningful contributions. First, it introduces a dynamic, time-resolved approach to sperm motility analysis, expanding beyond static single-timepoint metrics. Second, it demonstrates that intra-trajectory motility decay can be systematically quantified and is not merely attributable to random fluctuations. Third, it proposes a new analytical lens that may enrich future sperm quality assessments, particularly if future studies validate fatigue-like behavior with independent biochemical or functional endpoints.

Future research directions should prioritize experimental validation of the fatigue index by correlating motility decay patterns with mitochondrial activity, oxidative stress markers, and fertilization potential. Additionally, integrating real-time microenvironmental monitoring during sperm tracking may help disentangle intrinsic fatigue from external stressor effects. Longitudinal studies linking dynamic motility profiles with reproductive outcomes (e.g., IVF success rates) would further clarify the clinical significance of the fatigue phenomenon. Furthermore, sperm motility has been identified as a critical factor influencing the success rates of assisted reproductive technologies, including IVF, underscoring the clinical relevance of accurately assessing motility patterns (47–49). If future studies validate that motility decay directly contributes to impaired fertilization potential, the fatigue index could evolve into a clinically relevant dynamic biomarker for sperm quality assessment. Replication in independent datasets is essential to confirm the generalizability of the fatigue index, as current findings rely solely on a single ex vivo dataset.

As an exploratory study, the present work aims to generate new hypotheses regarding dynamic sperm motility deterioration rather than to confirm pre-specified clinical hypotheses.

In conclusion, this work establishes a conceptual foundation for sperm fatigue analysis and opens new avenues for dynamic phenotyping in andrology research. While preliminary, the findings highlight the potential value of integrating time-dependent motility metrics into sperm quality pipelines, emphasizing the need for cautious interpretation, independent validation, and thoughtful extension into clinical and translational contexts.

## 5. Conclusions

This study presents an exploratory framework for analyzing sperm motility deterioration through the concept of sperm fatigue, operationalized via a novel log-fold VSL index. Our findings demonstrate that intra-trajectory motility decline is a measurable, non-random phenomenon under ex vivo conditions, with biological plausibility grounded in mitochondrial dysfunction, oxidative stress, and ATP depletion mechanisms.

While limitations regarding causality, technical artifacts, and clinical applicability must be acknowledged, the systematic validation of fatigue patterns through multiple statistical approaches reinforces their potential biological relevance. Future studies integrating mechanistic assays and clinical outcomes will be critical to establish the role of sperm fatigue in reproductive success and to determine its value as a dynamic biomarker of sperm quality.

Overall, this work lays a conceptual foundation for the time-resolved phenotyping of sperm function, highlighting new opportunities for enhancing andrology research and clinical sperm assessment.

## 6. Acknowledgments

The authors would like to thank the developers and contributors of the VISEM-Tracking dataset for providing open access to high-quality sperm motility recordings, which made this study possible. The computational work was supported by open-source frameworks including scikit-learn, PyMC, matplotlib, and pandas. No external funding was received for the conduct or publication of this research.

## References

1. Leslie, S. W., Soon-Sutton, T. L., & Khan, M. A. B. (2024). Male infertility. In StatPearls [Internet]. Treasure Island (FL): StatPearls Publishing. Available from: https://www.ncbi.nlm.nih.gov/books/NBK562258/

2. Broekhuijse, M. L., Šoštarić, E., Feitsma, H., & Gadella, B. M. (2012). Application of computer-assisted semen analysis to explain variations in pig fertility. Journal of Animal Science, 90(3), 779–789. 10.2527/jas.2011-4311

3. Dcunha, R., Hussein, R. S., Ananda, H., Kumari, S., Adiga, S. K., Kannan, N., Zhao, Y., & Kalthur, G. (2022). Current insights and latest updates in sperm motility and associated applications in assisted reproduction. Reproductive Sciences, 29(1), 7–25. 10.1007/s43032-020-00408-y

4. Aghazarian, A., Huf, W., Pflüger, H., & Klatte, T. (2021). Standard semen parameters vs. sperm kinematics to predict sperm DNA damage. World Journal of Men’s Health, 39(1), 116–122. 10.5534/wjmh.190095

5. Villar, P. S., Vergara, C., & Bacigalupo, J. (2021). Energy sources that fuel metabolic processes in protruding finger-like organelles. FEBS Journal, 288(12), 3799–3812. 10.1111/febs.15620

6. Lassen, E., Pacey, A., Skytte, A. B., & Montgomerie, R. (2024). Recent decline in sperm motility among donor candidates at a sperm bank in Denmark. Human Reproduction, 39(8), 1618–1627. 10.1093/humrep/deae115

7. Chakraborty, S., & Saha, S. (2022). Understanding sperm motility mechanisms and the implication of sperm surface molecules in promoting motility. Middle East Fertility Society Journal, 27(4), Article 4. 10.1186/s43043-022-00094-7

8. Li, H., Shu, L., Yu, J., Xian, Z., Duan, H., Shu, Q., & Ye, J. (2021). Using Z-score to optimize population-specific DDH screening: A retrospective study in Hangzhou, China. BMC Musculoskeletal Disorders, 22(1), 344. 10.1186/s12891-021-04216-6

9. Gaffney, E. A., Ishimoto, K., & Walker, B. J. (2021). Modelling motility: The mathematics of spermatozoa. Frontiers in Cell and Developmental Biology, 9, 710825. 10.3389/fcell.2021.710825

10. Fernández-López, P., Garriga, J., Casas, I., Ramió-Lluch, L., Vilaseca, S., & Yeste, M. (2022). Predicting fertility from sperm motility landscapes. Communications Biology, 5, 1027. 10.1038/s42003-022-03954-0

11. Bae, J. W., Hwang, J. M., Lee, W. J., Kim, D. H., Yi, J. K., Ha, J. J., Oh, D. Y., & Kwon, W. S. (2024). Application of sperm motion kinematics and motility-related proteins for prediction of male fertility. Theriogenology, 218, 223–230. 10.1016/j.theriogenology.2024.02.007

12. Hackley, B., Kriebs, J. M., & Rousseau, M. E. (2007). Spermatozoon motility. In Mosby’s Guide to Women’s Health (2nd ed.). Elsevier Health Sciences.

13. Thambawita, V., Hicks, S. A., Storås, A. M., Nguyen, T., Andersen, J. M., Witczak, O., Haugen, T. B., Hammer, H. L., Halvorsen, P., & Riegler, M. A. (2023). VISEM-Tracking, a human spermatozoa tracking dataset. Scientific Data, 10(1), 260. 10.1038/s41597-023-02173-4

14. World Health Organization. (2021). WHO laboratory manual for the examination and processing of human semen (6th ed.). Available from: https://www.who.int/publications/i/item/9789240030787

15. Vasan, S. S. (2011). Semen analysis and sperm function tests: How much to test? Indian Journal of Urology, 27(1), 41–48. 10.4103/0970-1591.78424

16. Zhang, S., Bondarenko, O., Eskander Shazada, N., Cheng, Y., Roy Rahi, D., Rodina, M., et al. (2024). The plasma membrane contributes to temperature-induced spermatozoa motility kinetics in short-term stored sperm of common carp (*Cyprinus carpio*). Aquaculture, 583, 740588. 10.1016/j.aquaculture.2024.740588

17. Ishimoto, K., Gaffney, E. A., & Smith, D. J. (2017). Automatic tracking and motility analysis of human spermatozoa. IEEE Transactions on Biomedical Engineering, 64(10), 2381–2392. 10.1109/TBME.2017.2679301

18. Goodson, S. G., Zhang, Z., Tsuruta, J. K., Wang, W., & O’Brien, D. A. (2011). Classification of mouse sperm motility patterns using an automated multiclass support vector machines model. Biology of Reproduction, 84(6), 1207– 1215. 10.1095/biolreprod.110.088989

19. Tasoglu, S., Safaee, H., Zhang, X., Kingsley, J. L., Catalano, P. N., Gurkan, U. A., Nureddin, A., Kayaalp, E., Anchan, R. M., Maas, R. L., Tüzel, E., & Demirci, U. (2013). Exhaustion of racing sperm in nature-mimicking microfluidic channels during sorting. Small, 9(20), 3374–3384. 10.1002/smll.201300020

20. Wu, H., Yu, X., Wang, Q., Zeng, Q., Chen, Y., Lv, J., Wu, Y., Zhou, H., Zhang, H., Liu, M., Zheng, M., Zhao, Q., Guo, P., Feng, W., Zhang, X., & Tian, L. (2021). Beyond the mean: Quantile regression to differentiate the distributional effects of ambient PM2.5 constituents on sperm quality among men. Chemosphere, 263, 131496. 10.1016/j.chemosphere.2021.131496

21. Keihani, S., Verrilli, L. E., Zhang, C., Presson, A. P., Hanson, H. A., Pastuszak, A. W., Johnstone, E. B., & Hotaling, J. M. (2021). Semen parameter thresholds and time-to-conception in subfertile couples: How high is high enough? Human Reproduction, 36(8), 2121–2133. 10.1093/humrep/deab133

22. Li, H., Shu, L., Yu, J., Xian, Z., Duan, H., Shu, Q., & Ye, J. (2021). Using Z-score to optimize population-specific DDH screening: A retrospective study in Hangzhou, China. BMC Musculoskeletal Disorders, 22(1), 344. 10.1186/s12891-021-04216-6

23. Funke, M., Yang, Y., Lahtinen, A., Benninghoven-Frey, K., Kliesch, S., Neuhaus, N., Stukenborg, J. B., & Jahnukainen, K. (2021). Z-scores for comparative analyses of spermatogonial numbers throughout human development. Fertility and Sterility, 116(3), 713–720. 10.1016/j.fertnstert.2021.04.019

24. Tanaka, T., Kojo, K., Nagumo, Y., Ikeda, A., Shimizu, T., Fujimoto, S., Kakinuma, T., Uchida, M., Kimura, T., Kandori, S., & Nishiyama, H. (2024). A new clustering model based on the seminal plasma/serum ratios of multiple trace element concentrations in male patients with subfertility. Reproductive Medicine and Biology, 23(1), e12584. 10.1002/rmb2.12584

25. Hicks, S. A., Andersen, J. M., Witczak, O., Thambawita, V., Halvorsen, P., Hammer, H. L., Haugen, T. B., & Riegler, M. A. (2019). Machine learning-based analysis of sperm videos and participant data for male fertility prediction. Scientific Reports, 9(1), 16770. 10.1038/s41598-019-53217-y

26. Naik, N., Roth, B., & Lundy, S. D. (2024). Artificial intelligence for clinical management of male infertility: A scoping review. Current Urology Reports, 26(1), 17. 10.1007/s11934-024-01239-z

27. Paffoni, A., Somigliana, E., Boeri, L., & Viganò, P. (2022). The statistical foundation of the reference population for semen analysis included in the sixth edition of the WHO manual: A critical reappraisal of the evidence. Human Reproduction, 37(10), 2237–2245. 10.1093/humrep/deac161

28. Luck, S. (2020). Nonoverlap proportion and the representation of point-biserial variation. PLoS One, 15(12), e0244517. 10.1371/journal.pone.0244517

29. Ghuman, N. K., Shukla, K. K., Nandagopal, S., Raikar, S., Kumar, S., Kathuria, P., Choudhary, D., Elhence, P., & Singh, P. (2023). Explaining the unexplained: Examining the predictive value of semen parameters, sperm DNA fragmentation and metal levels in unexplained infertility. Journal of Human Reproductive Sciences, 16(4), 317–323. 10.4103/jhrs.jhrs_140_23

30. Nichols, K., & Holmes, A. (2007). Non-parametric procedures. In K. Friston, J. Ashburner, S. Kiebel, T. Nichols, & W. Penny (Eds.), Statistical parametric mapping: The analysis of functional brain images (pp. 253–272). Academic Press. 10.1016/B978-012372560-8/50022-0

31. Emmert-Streib, F. (2024). Trends in null hypothesis significance testing: Still going strong. Heliyon, 10(21), e40133. 10.1016/j.heliyon.2024.e40133

32. Pernet, C. (2015). Null hypothesis significance testing: A short tutorial. F1000Research, 4, 621. 10.12688/f1000research.6963.3

33. Efron, B., & Narasimhan, B. (2020). The automatic construction of bootstrap confidence intervals. Journal of Computational and Graphical Statistics, 29(3), 608–619. 10.1080/10618600.2020.1714633

34. Xu, Z., Merino-Sanjuan, M., Mangas-Sanjuan, V., & García-Arieta, A. (2021). Estimators and confidence intervals of f2 using bootstrap methodology for the comparison of dissolution profiles. Computer Methods and Programs in Biomedicine, 212, 106449. 10.1016/j.cmpb.2021.106449

35. van Zyl, C., Ye, X., & Naidoo, R. (2024). Harnessing explainable artificial intelligence for feature selection in time series energy forecasting: A comparative analysis of Grad-CAM and SHAP. Applied Energy, 353(Part A), 122079. 10.1016/j.apenergy.2023.122079

36. Patel, A., Baek, S., & Keutzer, K. (2024). Interpretable machine learning for bioengineering applications: From global to local explanations. Advanced Intelligent Systems, 6(5), 2400304. 10.1002/aisy.202400304

37. Huang, X., & Marques-Silva, J. (2024). On the failings of Shapley values for explainability. International Journal of Approximate Reasoning, 171, 109112. 10.1016/j.ijar.2023.109112

38. Wang, X., Niu, Y., & Li, X. (2017). Bayesian ROC curve estimation under binormality using an ordinal category likelihood. Communications in Statistics -Theory and Methods, 47(18), 4628–4640. 10.1080/03610926.2017.1380830

39. Durairajanayagam, D., Singh, D., Agarwal, A., & Henkel, R. (2021). Causes and consequences of sperm mitochondrial dysfunction. Andrologia, 53(1), e13666. 10.1111/and.13666

40. Vahedi Raad, M., Firouzabadi, A. M., Tofighi Niaki, M., Henkel, R., & Fesahat, F. (2024). The impact of mitochondrial impairments on sperm function and male fertility: A systematic review. Reproductive Biology and Endocrinology, 22(1), 83. 10.1186/s12958-024-01252-4

41. Mai, Z., Yang, D., Wang, D., Zhang, J., Zhou, Q., Han, B., & Sun, Z. (2024). A narrative review of mitochondrial dysfunction and male infertility. Translational Andrology and Urology, 13(9), 2134–2145. 10.21037/tau-24-262

42. Tomlinson, M. J., Pooley, K., Simpson, T., Newton, T., Hopkisson, J., Jayaprakasan, K., Jayaprakasan, R., Naeem, A., & Pridmore, T. (2010). Validation of a novel computer-assisted sperm analysis (CASA) system using multitarget-tracking algorithms. Fertility and Sterility, 93(6), 1911–1920. 10.1016/j.fertnstert.2008.12.064

43. Canonico, L. F., De Clemente, C., Fardilha, M., Ferreira, A. F., Maremonti, M. I., Dannhauser, D., Causa, F., & Netti, P. A. (2024). Exploring altered bovine sperm trajectories by sperm tracking in unconfined conditions. Frontiers in Veterinary Science, 11, 1358440. 10.3389/fvets.2024.1358440

44. Morbeck, D. E., Leonard, P. H., Weaver, A. L., Shimek, K. M., Bouwsma, E. V., & Coddington, C. C. (2011). Sperm morphology: Classification drift over time and clinical implications. Fertility and Sterility, 96(6), 1350–1354. 10.1016/j.fertnstert.2011.08.036

45. Appell, R. A., & Evans, P. R. (1977). The effect of temperature on sperm motility and viability. Fertility and Sterility, 28(12), 1329–1332. 10.1016/s0015-0282(16)42978-x

46. Hoang-Thi, A. P., Dang-Thi, A. T., Phan-Van, S., Nguyen-Ba, T., Truong-Thi, P. L., Le-Minh, T., Nguyen-Vu, Q. H., & Nguyen-Thanh, T. (2022). The impact of high ambient temperature on human sperm parameters: A meta-analysis. Iranian Journal of Public Health, 51(4), 710–723. 10.18502/ijph.v51i4.9232

47. Cheng, Q., Li, L., Jiang, M., Liu, B., Xian, Y., Liu, S., Liu, X., Zhao, W., & Li, F. (2022). Extend the survival of human sperm in vitro in non-freezing conditions: Damage mechanisms, preservation technologies, and clinical applications. Cells, 11(18), 2845. 10.3390/cells11182845

48. Dhumal, S. S., Naik, P., Dakshinamurthy, S., & Sullia, K. (2021). Semen pH and its correlation with motility and count: A study in subfertile men. JBRA Assisted Reproduction, 25(2), 172–175. 10.5935/1518-0557.20200080

49. Dcunha, R., Hussein, R. S., Ananda, H., Kumari, S., Adiga, S. K., Kannan, N., Zhao, Y., & Kalthur, G. (2022). Current insights and latest updates in sperm motility and associated applications in assisted reproduction. Reproductive Sciences, 29(1), 7–25. 10.1007/s43032-020-00408-y

